# Molecular Studies of Phages-*Klebsiella pneumoniae* in a Mucoid Environment: Innovative use of mucolytic agents prior to the administration of lytic phages

**DOI:** 10.1101/2023.08.02.551690

**Authors:** Olga Pacios, Lucía Blasco, Concha Ortiz-Cartagena, Inés Bleriot, Laura Fernández-García, María López, Antonio Barrio-Pujante, Álvaro Pascual, Felipe Fernández Cuenca, Luis Martínez-Martínez, Belén Aracil, Jesús Oteo-Iglesias, María Tomás

## Abstract

Mucins are important glycoproteins that form a protective layer throughout the gastrointestinal and respiratory tracts. There is scientific evidence of increase in phage-resistance in the presence of mucin for some bacterial pathogens. Manipulation in mucin composition may ultimately influence the effectiveness of phage therapy. In this work, two clinical strains of *K. pneumoniae* (K3574 and K3325), were exposed to the lytic bacteriophage vB_KpnS-VAC35 in the presence and absence of mucin on a long-term co-evolution assay, in an attempt to mimic *in vitro* the exposure to mucins that bacteria and their phages face *in vivo*. Enumerations of the bacterial and phage counts at regular time intervals were conducted, and extraction of the genomic DNA of co- evolved bacteria to the phage, the mucin and both was performed. We determined the frequency of phage-resistant mutants in the presence and absence of mucin and including a mucolytic agent (N-acetyl L-cysteine, NAC), and sequenced these conditions using Nanopore. We phenotypically demonstrated that the presence of mucin induces the emergence of bacterial resistance against lytic phages, effectively decreased in the presence of NAC. In addition, the genomic analysis revealed some of the genes relevant to the development of phage resistance in long-term co- evolution, with a special focus on the mucoid environment. Genes involved in the metabolism of carbohydrates were mutated in the presence of mucin. In conclusion, the use of mucolytic agents prior to the administration of lytic phages could be an interesting therapeutic option when addressing *K. pneumoniae* infections in environments where mucin is overproduced.

## Introduction

*Klebsiella pneumoniae* is a Gram-negative opportunistic pathogen that causes urinary tract, wound and soft tissue infections, pneumonia, and even life-threatening sepsis (1, 2). Moreover, the recent increase of carbapenemase-producing strains of *K. pneumoniae* worldwide, together with its ability to grow in biofilm and to acquire plasmids conferring antibiotic (multi)resistance, underlie the importance of developing innovative and effective strategies against *K. pneumoniae* infections (3, 4).

In this context, the use of bacteriophages (or phages), viruses that specifically target bacteria in a highly effective and safe manner, is being evaluated as a therapeutic approach against bacterial infections, especially antibiotic-resistant ones (4, 5). Nevertheless, just as it happens with antibiotics, the emergence of phage-resistant mutants is a major hurdle to the establishment of phage therapy (4, 6). Indeed, to counter phage infection, bacteria display several defence mechanisms: mutation of the receptor recognized by a particular phage to inhibit adsorption (surface mutation) (7), induction of programmed death cell, known as abortive infection (Abi) (8), translation of nucleases that specifically degrade the phage DNA (CRISPR-Cas, restriction- modification… (9, 10)), etc. Despite the inconvenience of resistant bacteria against phages, their compassionate use in clinics has been approved in many countries and has already saved many life-threatening infectious patients (11–13).

Cystic Fibrosis (CF), an autosomal recessive genetic disorder that produces mutations in the cystic fibrosis transmembrane conductance regulator (CFTR) protein, is characterized by an overproduction of viscous mucins, since lack of CFTR function reduces airway mucus fluidity and influences hydration and mucin viscosity in the airways (14). This allows the trapping of inhaled bacteria in the lungs and explains why CF patients often become colonized by pathogens from an early age, which can lead to chronic infections (15). Even if a few typical bacteria are traditionally involved in CF lung infections, such as *Staphylococcus aureus* and *Pseudomonas aeruginosa*, CF patients are susceptible to infection by other opportunistic pathogens, including *K. pneumoniae* (16, 17).

To improve therapeutic outcomes in phage therapy, the arising of phage-resistant bacteria in the complex *in vivo* context needs to be exploited. Furthermore, not many studies address the efficiency of phage in long-term evolutionary experiments, nor look at phage co-evolution during phage treatments, as reviewed by Mouloton-Brown in 2018 (18).

One of the main components of the gastrointestinal and respiratory tracts are mucins. Mucins are high-molecular-weight proteins that are glycosylated and can be transmembrane (forming a protective “brush” border on the epithelium) or gel-forming (providing hydration and protection from shear stress) (19, 20). They protect the intestinal mucosa from physical contact with commensal bacteria, as well as from invasion of intruders and pathogens (21) . Changes in mucin expression are relevant in inflammatory and neoplastic disorders of the gastrointestinal tract, being important in the etiology of some infectious diseases, such as *Helicobacter pylori* gastritis (22).

In the present work, we have used two bacteriemic-causing clinical isolates of *K. pneumoniae*, named K3574 and K3325, and exposed them to the lytic bacteriophage vB_KpnS-VAC35 in the presence and absence of mucin on a long-term co-evolution assay, intending to study the phage resistance in a mucoid environment. We determined the relationship between mucin and the difficulties in applying phage therapy, and we included a mucolytic agent to improve the use of phages by avoiding the emergence of resistance.

## Results

### Infectivity of phage vB_KpnS-VAC35

The two strains exhibiting the highest efficiency of plating (EOP) values (K3574 and K3325) were chosen for further assays (Figure 1a). Optical density growth curves showed good lytic activity of vB_KpnS-VAC35 in these strains at a multiplicity of infection (MOI) of 0.1 (purple line) and 1 (orange line) (Figure 1 b) and c), respectively). The isolate K3325 was less well infected by vB_KpnS-VAC35 than the isolation host, K3574.

**Figure 1:**
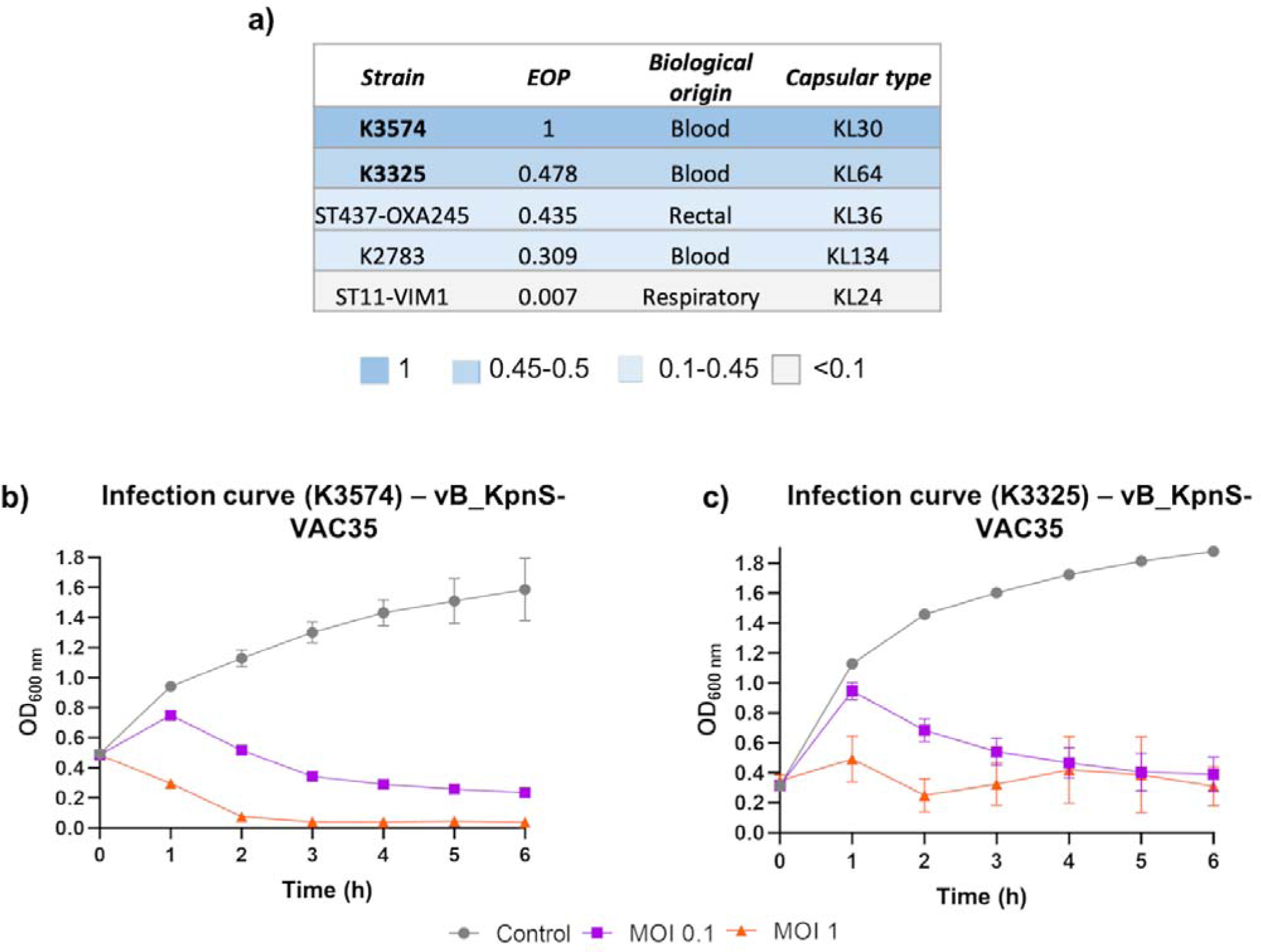
a) EOP of vB_KpnS-VAC35 on some clinical isolates of *K. pneumoniae*, with different capsular type. b) and c) Infection curves of clinical isolates K3574 and K3325 by the lytic phage vB_KpnS-VAC35, prior to co-evolution, at MOI 0.1 and 1.

### Co-evolution of K3574 and K3325 with the phage vB_KpnS-VAC35 in a mucoid environment

For the co-evolution experiment, the initial bacterial inoculum was 2x10^8^ colonies forming units per mL (CFU/mL) for K3574 and 10^8^ CFU/mL for K3325 (Figure 2). As we infected both cultures in the exponential growth phase at a MOI=1, the initial phage concentration was 2x10^8^ plaque forming units per mL (PFU/mL) and 10^8^ PFU/mL, respectively (Figure 2). The CFU counts remained stable at around 10^10^ CFU/mL for both strains at every condition tested, whereas the PFUs fluctuated slightly more: in what concerns the isolate K3574 co-adapted to the phage, PFU counts ranged from 10^9^ PFU/mL at 1 day post-infection (dpi) to 10^7^ PFU/mL at 15 dpi, whereas in the presence of mucin these counts reached 10^8^ PFU/mL at 9 dpi, then slightly decreased till 10^7^ PFU/mL at 15 dpi (Figure 2a), which corresponds to the 1:100 dilution performed every day along the experiment. Regarding the isolate K3325, co-evolution lasted 6 days as the PFU numbers dropped to 0 in the absence of mucin (Figure 2b). Nonetheless, in the presence of this compound, 10^3^ PFU/mL of vB_KpnS-VAC35 were assessed at 6 dpi.

**Figure 2:**
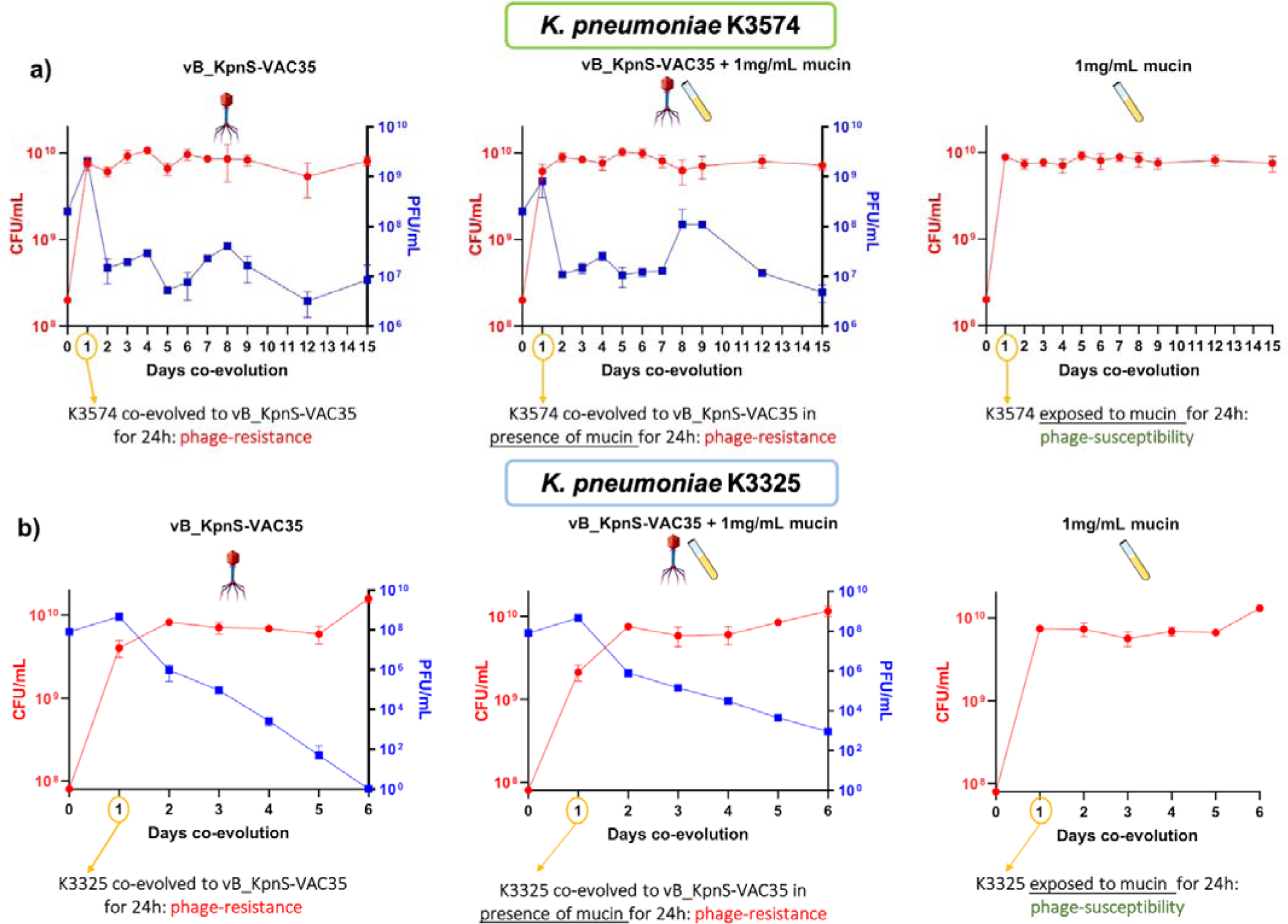
Titration of the colony forming units (CFU, left Y-axis) and plaque forming units (PFU, right Y-axis) per mL during the co-evolution experiments between clinical isolates K3574 (a)) and K3325 (b)) and the lytic bacteriophage vB_KpnS-VAC35, in the absence and presence of mucin (1 mg/mL).

### Assessment of phage-resistance

#### a. Spot test

With the aim to assess the effect that the presence of mucin in the media will have on the bacterial susceptibility to the adapted phages, a spot test of *K. pneumoniae* K3574 and K3325 prior to the co-evolution (named as “WT” in Figure 3), but also co-evolved in the presence of mucin, the phage, and both during 15 and 6 days, respectively, was conducted. As expected, WT and mucin-adapted cells (K3574_ad15_m and K3325_ad6_m) were the only conditions in which phage-susceptibility was kept (Figure 3). However, we observed differences when comparing the infection established by the non-adapted phage (vB_KpnS-VAC35_WT) and the adapted ones (vB_KpnS-VAC35_ad15 and vB_KpnS-VAC35_ad15_m), which were isolated after 15 days of co-evolution with K3574 and produced more turbid spots (Figure 3, middle and right columns). The presence of more colonies growing inside the lytic halos of vB_KpnS-VAC35_ad15_m compared to the infection established by vB_KpnS-VAC35_ad15 suggested that mucin either impaired the ability of the phage to lyse, or it enhanced the bacterial defence to the phage (as it has already been documented in the literature for different microorganisms (23, 24)).

**Figure 3:**
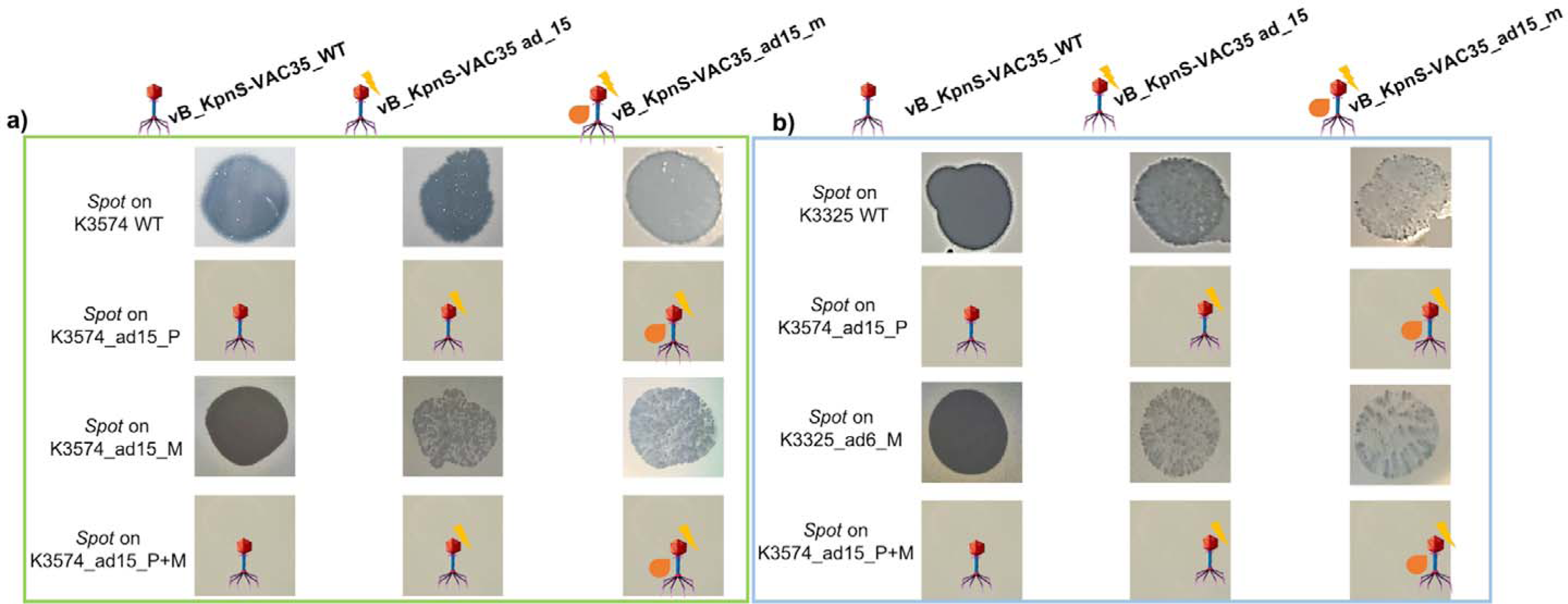
Spot tests of phages prior to co-evolution (vB_KpnS-VAC35 WT), co-evolved to K3574 in the absence and in the presence of mucin (vB_KpnS-VAC35_ad15 and vB_KpnS-VAC35_ad15_m, respectively). A) Spot test using the clinical isolate *K. pneumoniae* K3574 WT, adapted 15 days to the phage, to mucin and to both. B) Spot test using the clinical isolate *K. pneumoniae* K3325 WT, adapted 6 days to the phage, to mucin and to both.

#### b. Frequency of arising of resistant mutants in the presence of the mucolytic N-acetyl L-cysteine (NAC)

To quantitatively assess the effect that the presence of mucin had during bacteria and phage co-evolution, we determined the frequency of phage-resistant mutants to vB_KpnS-VAC35. A condition in which *K. pneumoniae* clinical isolates K3574 and K3325 were incubated in presence of NAC for 15 and 6 days, respectively, was included. Consistently with the infection curves of vB_KpnS-VAC35 in these two strains, we obtained a higher frequency for K3325 than K3574 (Figure 4, orange bar). In the case of phage-exposed bacteria, either in the presence of mucin, NAC, or only the phage, no statistical difference was observed (Figure 4). Importantly, cells exposed to NAC displayed a statistically significant reduction in this frequency compared to the cells exposed to mucin.

**Figure 4:**
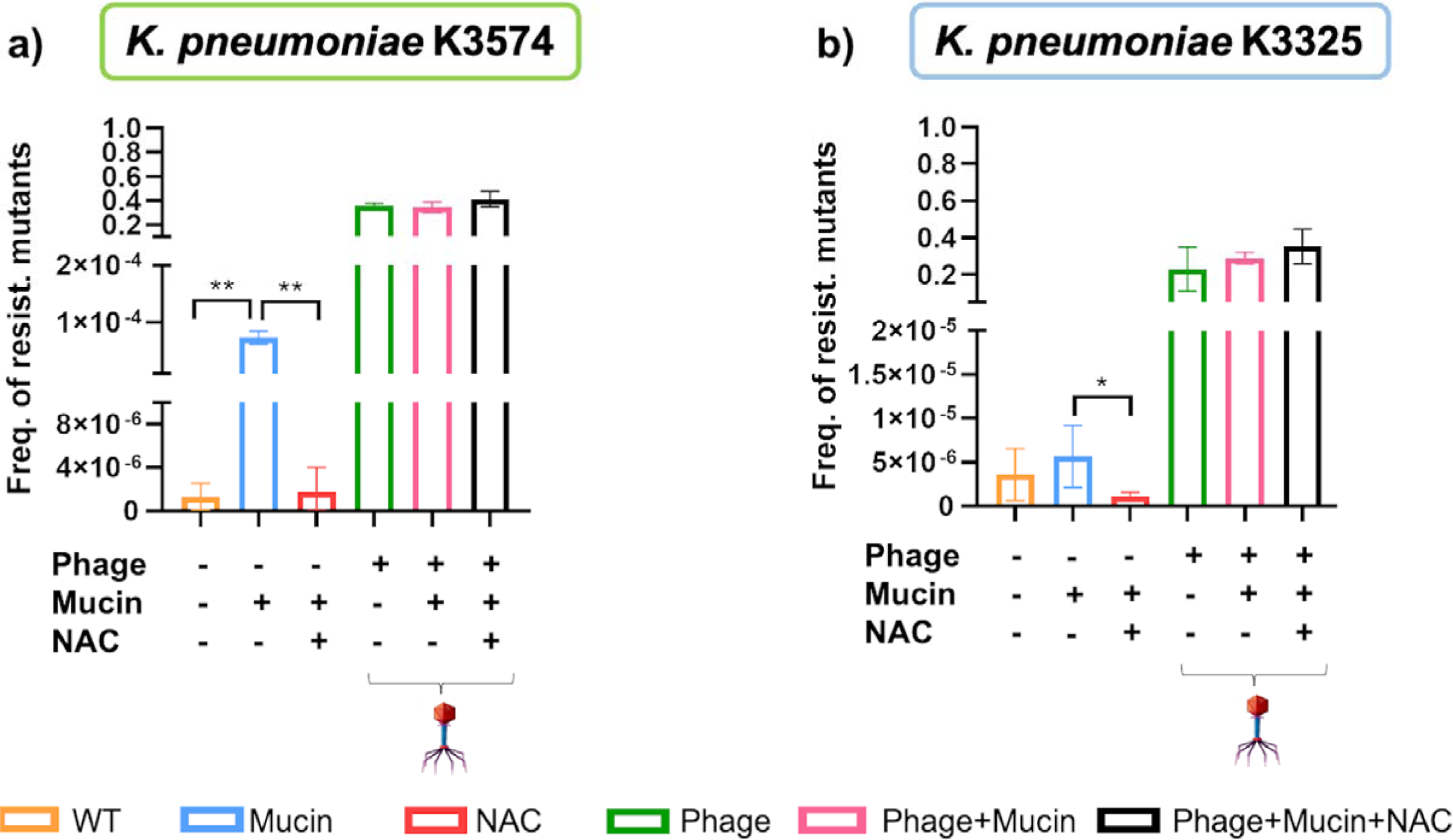
Frequency of occurrence of resistant mutants for *K. pneumoniae* K3574 (a) and *K. pneumoniae* K3325 (b). The statistical analysis (t-test) was performed with GraphPad Prism v.9. **: p-value < 0.001; *: p-value 0.042. Absence of asterisk corresponds to non-statistical significance.

### Genomic analysis of *K. pneumoniae* strains before and after the co-evolution in the presence of mucin

Genomic analysis of the clinical strain K3574 adapted to mucin, to the phage alone or to both agents, were performed.

The reference genome of this strain (BioSample code SAMEA3649560, European BioProject PRJEB10018) possesses 5,635,279 pb (5561 coding sequences) with a GC content of 57.1%, a sequence-type ST3647 and a capsular type KL30. We extracted the bacterial DNA at 15 dpi of this isolate co-evolved to the phage (K3574_ad15_P), to mucin (K3574_ad15_M) and to both (K3574_ad15_P+M).

A Venn diagram was used to visualize the different and overlapping protein clusters displayed by the four complete genomes taken into consideration (Figure 5 b). In total, 5000 common protein clusters were found between the strain prior to co-evolution and bacteria co-evolved to the phage (K3574_ad15_P), whereas 4938 were found between K3574_WT and the mucin-adapted cells (K3574_ad15_M), and 4947 common protein clusters between K3574_WT and cells co-evolved to the phage in the presence of mucin (K3574_ad15_P+M). No specific protein clusters were found for the strain adapted only to mucin (K3574_ad15_M) and to phage and mucin together (K3574_ad15_P+M), while 3 unique protein clusters were found in the isolate co-evolved to the phage alone (K3574_ad15_P) (Figure 5 b).

**Figure 5:**
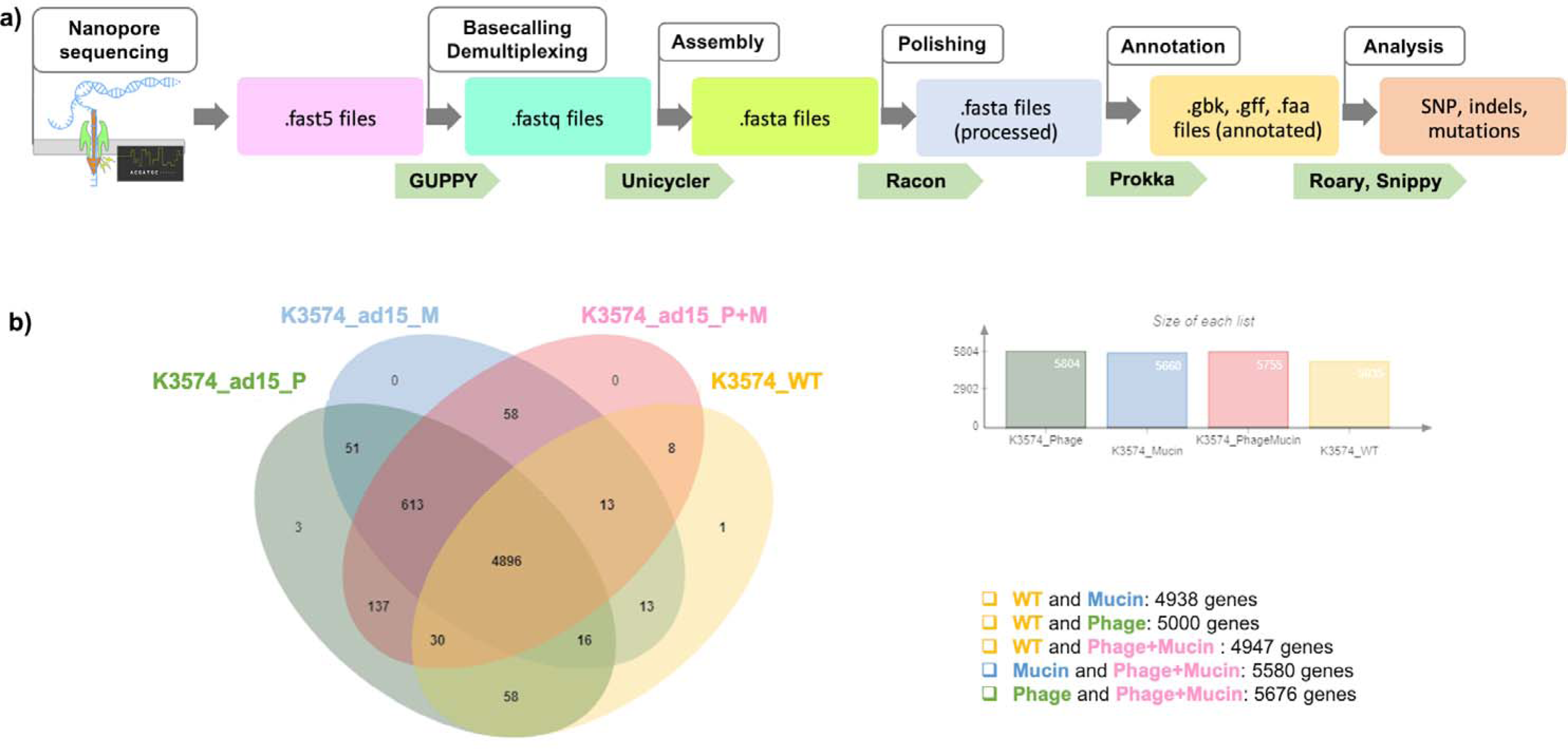
a) Schematic representation of the workflow followed from the whole-genome sequencing with Nanopore till the analyse of the genomic sequences. B) Venn diagram performed with OrthoVenn2 to visualize the overlapped and unique protein clusters in the four complete genomes analysed.

Comparison of the genomes revealed important mutations (Figure 6, Table 1). Among the genes in which nucleotide changes were found, we highlight several interesting ones grouped into different categories: concerning the bacterial defense mechanisms to phage infection, a tRNA- guanosine (18)-2’-O-methyltransferase carrying a nucleotide deletion in the position 362 was found in the case of K3574_ad15_P and K3574_ad15_P+M, whereas the non-infected isolate (K3574_ad15_M) had the intact locus compared to the K3574_WT. This change (c.-362G) leads to a frameshift mutation translated into two truncated versions of the methyltransferase. Furthermore, the antitoxin HigA displayed mutations in the phage-infected cultures (K3574_ad15_P and K3574_ad15_P+M) that led to two different truncated proteins, whereas the non-infected strain had fewer changes that led to a shorter, unique version. Furthermore, the autoinducer 2-binding protein LsrB presented the same nucleotide deletion in the strains that co-evolved to the phage, also corresponding to a frameshift mutation (therefore a truncated protein lacking 11 amino acids); this was absent in the isolate exposed only to mucin.

**Figure 6:**
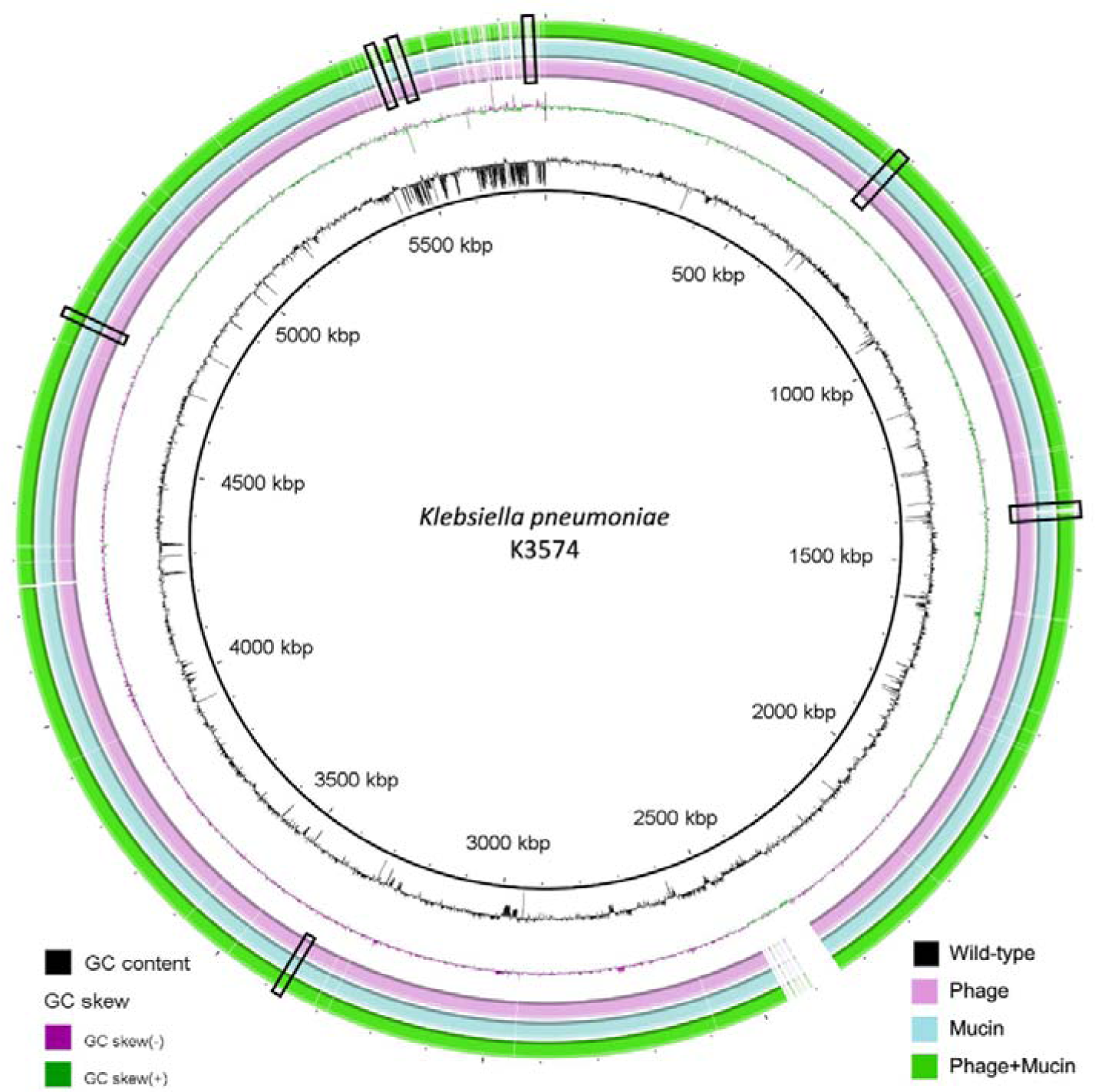
Comparative genomic analysis of *K. pneumoniae* K3574 co-evolved to phage alone (pink ring), mucin alone (blue ring) and both (green ring) constructed with the BLAST Ring Image Generator (BRIG). The sequence corresponding to K3574_WT is located on the innermost side (black ring). The double ring adjacent to the reference sequence represents the GC content (black) and the GC skew (dark purple and dark green). The white parts of the rings represent absent or divergent content and are squared in black.

**Table 1:**
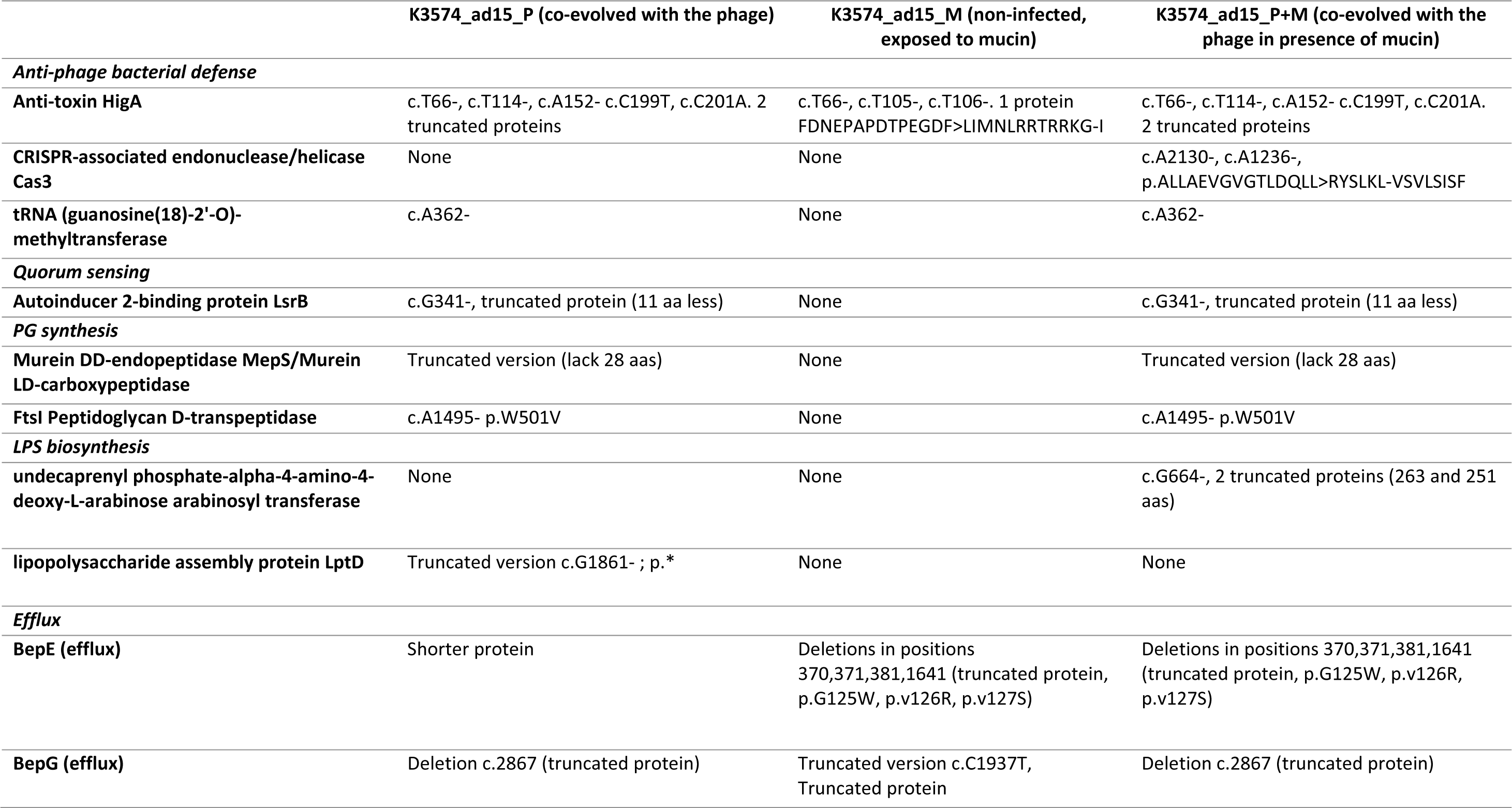
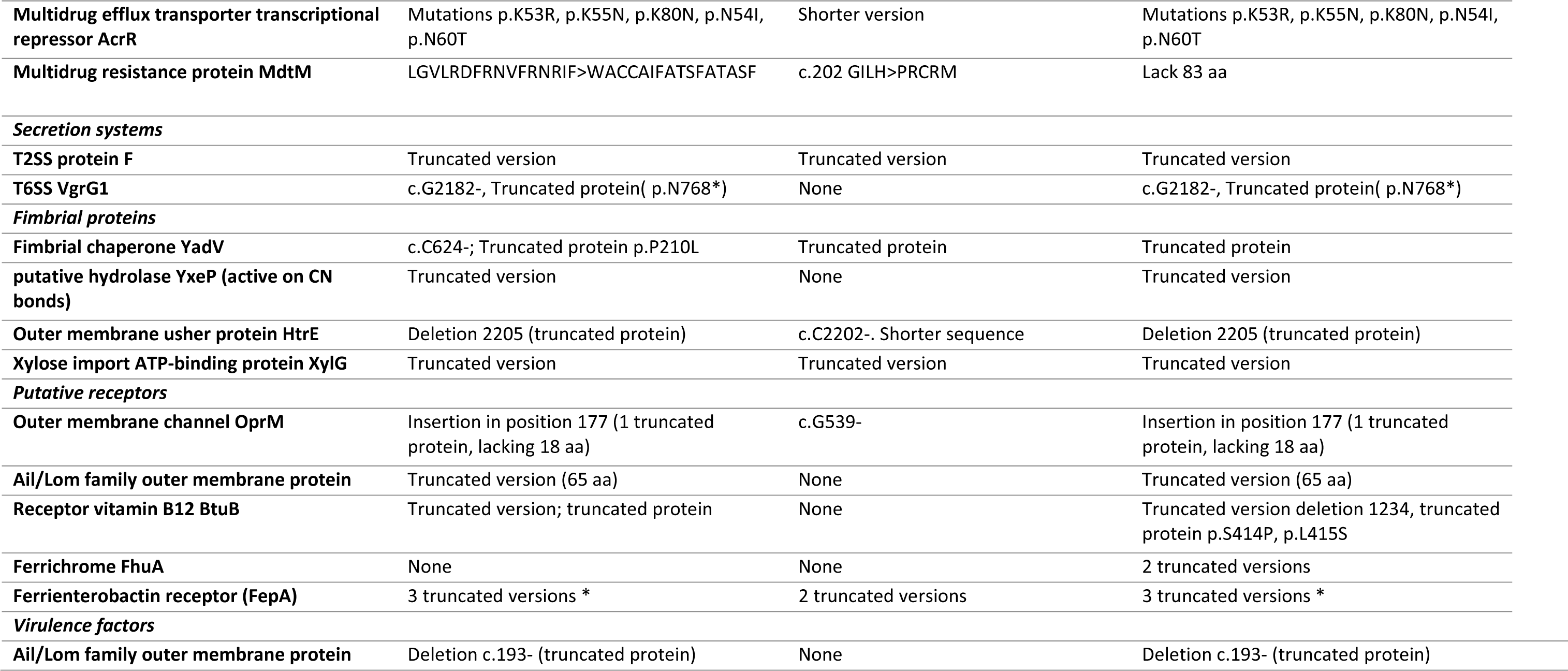
Mutations found in the four complete genomes analyzed, grouped into different categories by function. “c.” corresponds to the nucleotide changes found in the coding sequences, while “p.” stands for protein sequences. Asterisks indicate that mutations are the same between the studied conditions.

Mutations in the core gene involved in Type VI secretion system, *vgrG,* were found in these same two strains but absent in the strain adapted only to mucin (K3674_ad15_M). Similarly, the same changes were found in the coding sequence of the outer membrane protein Ail (*attachment invasion locus*) of K3574_ad15_P and K3574_ad15_P+M, being intact in the isolate K3574_ad15_M. Interestingly, *fhuA*, the gene encoding a ferrichrome transporter protein, showed a frameshift mutation that led to two truncated versions of this protein in K3574_ad15_P+M, whereas this mutation was absent in K3574_ad15_P and K3574_ad15_M.

Mutations were found in genes involved in the metabolism of carbohydrates only for the strains adapted 15 days to mucin alone and to the phage in presence of mucin. For instance, nucleotide changes were observed in the gene *licC*, involved in the phosphoenolpyruvate-dependent sugar phosphotransferase system (PTS), in *rbsA, rbsB* and *fruK*, encoding the ribose and fructose import ATP-binding proteins RbsA/RbsB and FruK, respectively, or in *malI,* encoding a maltose regulon regulatory protein. More nucleotide changes in other relevant genes are reported in Table 1.

## Discussion

Most of the works studying the interactions between pathogenic bacteria and their phages are generally carried out in well-defined laboratory conditions. However, these microbes in a natural environment develop complex interactions on mucosal surfaces of the vertebrate host (23). In 2022, de Freitas *et al.* demonstrated that co-evolution of *Flavobacterium columnare* to its virulent phage V156 in presence of mucin dramatically increased the acquisition of spacers in the CRISPR arrays of *F. columnare* (*23*), thus increasing immunity to this phage. This work highlights the need to consider both biotic and abiotic variables if bacteriophages are to be used therapeutically. It is thus essential to take a close look at the study of how mucins and other mucosal components influence the acquisition of bacterial resistance towards lytic phages, so that the therapeutic potential of these could be better understood in the *in vivo* system.

Along the co-evolution performed with *K. pneumoniae* isolates K3574 and K3325, we observed that phage mutants arose at the first day post-infection (Figure 2), similar to what has been claimed in previous co-evolution works (25). Interestingly, both phage production (from 1 dpi onwards) and evolved *K. pneumoniae* populations seemed to stabilize over the days, consistently with other studies (26). This is likely due to the fact that vB_KpnS-VAC35 selects for resistant bacterial mutants. Due to its gelatinous nature, mucin limits the diffusion of bacteria through this space and facilitates the interaction of phages with bacteria, a less well-studied function but already documented in literature (23, 27). In 2013, Barr *et al.* proposed a model in which bacteriophages would bind to mucins using the Ig-like domains present in many structural proteins, concentrate there, and protect humans and other metazoans against bacterial invaders (27, 28). Taken together, these two reasons could explain why the phage counts were higher in the presence of mucin than in the absence of this compound (Figure 2).

Interestingly, when the lytic phage was co-evolved to K3574 in the mucoid environment (vB_KpnS-VAC35_ad15_m), it showed an impaired infection ability compared to the adapted phage in the absence of mucin (vB_KpnS-VAC35_ad15) (Figure 3). As expected, no spot was visible for the cells exposed to the phage either in the presence or the absence of mucin, leading to the conclusion that resistance arose as a result of the co-evolution process. We phenotypically confirmed that mucin increased the frequency of resistant mutants for *K. pneumoniae* K3574 and K3325 strains. Furthermore, NAC effectively reduced this frequency in the case of the cells incubated with this mucolytic (Figure 4). This resistant phenotype could be due to modification of the phage receptor; however, as this strategy represents an important fitness cost for bacteria, these have developed other strategies to avoid phage attachment (29). For instance, receptors can be masked, preventing recognition while retaining function. Capsules or exopolysaccharides provide phage resistance in *Staphylococcus spp.* (*30*), *Pseudomonas spp.* and *K. pneumoniae* (31), and these bacterial structures can be favorized in the presence of mucosal components such as mucins.

The analysis of protein clusters suggested that the presence of the phage in this long-term co- evolution experiment was the main driver in the acquisition of mutations.The genomic analysis of *K. pneumoniae* K3574 adapted to the phage alone and in presence of mucin revealed mutations in some proteins involved in the bacterial defense to phages, such as methyltransferases, the HigA antitoxin, the *quorum sensing* autoinducer LsrB or the type 6 secretion system VgrG, as reported in other works (32, 33) (Figure 6). In the presence of mucin, mutations were observed in the genes encoding proteins that were involved in the carbohydrates metabolism, such as in the PTS system, which is a major carbohydrate active-transport system that catalyzes the phosphorylation of incoming sugar substrates concomitant with their translocation across the cell membrane (34). Since changes concerning the synthesis, secretion or structure of mucins have been linked to gastrointestinal and respiratory disorders, manipulation of mucin may ultimately influence the microbiota and the effectiveness of phage therapy for bacterial imbalances (21), and the use of a mucolytic agent as an adjuvant of lytic phages could be an interesting therapeutic option to take into consideration. It has been shown that the presence of bacteria upregulates mucin production and enhances their encapsulation by mucin in the colon, so this could be even more important in CF patients in which overproduction of mucins leads to lung chronic infections (35). Importantly, the trade-off costs that phage pressure and co-evolution represent for bacteria might render them less virulent in case of mutations in surface virulence factors, so maximizing the fitness costs that come with co-evolution may ultimately enhance the long-term efficacy of phage therapy. Optimization of these fitness costs could be a relevant factor to enhance the patient’s prognosis (36).

All in all, this study sheds some light in the phage resistance behavior that might be expected for some clinical strains of *K. pneumoniae* in a mucoid environment, and takes a deeper look at the increase resistance that mucins induce to phages, already reported in literature (23). Evolutionary dynamics between bacterial pathogens and their natural predators in *in vivo* environments where mucin overproduction occurs deserve further investigation, which could help clinicians to predict the success of a particular phage administered to counteract infections. Finally, our results showed an innovative to option could be the application of mucolytic agents prior to the administration of lytic phages against by *K. pneumoniae* infections environments where mucin is overproduced as in cystic fibrosis disease. However, would be necessary to carry out more studies that include broad number clinical isolates to confirm this innovative therapeutic option.

## Materials and methods

### Bacterial strains and growth conditions

*K. pneumoniae* clinical strains K3574 and K3325 came from the National Centre for Microbiology (Carlos III Health Institute, Spain) previously analyzed^23^. All the bacterial strains were cultivated using Luria-Bertani broth (LB, 1% tryptone, 0.5% yeast extract and 0.5% NaCl). When required, purified mucin from porcine stomach (SigmaAldrich®), previously diluted in distilled water and autoclave-sterilized, was added at a final concentration of 1 mg/mL. NAC was also purchased from SigmaAldrich®, diluted with nuclease-free water, filter-sterilized and added to a final concentration of 10 mM.

### Establishment of the infectivity of the phage

#### a. Efficiency of plating (EOP)

The EOP assay was done as previously described by Kutter *et al.* ^24^, calculated as the ratio between the phage titre (PFU/mL) in the test strain and the titre in the isolation host (*K. pneumoniae* K3574). For both assays, TA-soft medium (1% tryptone, 0.5% NaCl and 0.4% agar) was used to make plates by the top-agar method^25^. Strains exhibiting susceptibility to phage infection in the spot test performed by Bleriot *et al.* in a previous work ^23^ were selected for the EOP assay.

#### b. Infection curves

To assess the lytic capacity of vB_KpnS-VAC35^23^, infection curves at different MOI were performed. Overnight cultures of the clinical isolates of *K. pneumoniae* K3574 and K3325 were diluted 1:100 in LB broth and then incubated at 37°C at 180 rpm until an early exponential phase (OD_600 nm_ = 0.3- 0.4) was reached. Then, vB_KpnS-VAC35 was added to the cultures at MOI of 0.1 and 1, and OD_600 nm_ was measured during 6 hours at 1-hour intervals.

### Co-evolution between vB_KpnS-VAC35 and *K. pneumoniae* strains K3574 and K3325

The bacterial strains were incubated in 20 mL LB-containing flasks at 37°C and 180 rpm for 6 (K3325) or 15 days (K3574); the flasks were infected with vB_KpnS-VAC35 at a MOI=1 in the presence and absence of porcine mucin at a final concentration of 1 mg/mL, and a non-infected control of the bacterial isolate growing in presence of 1 mg/mL mucin was included. The infections with the phage were performed at OD≈of 0.4. From this moment and every 24 hours, each condition was 1:100 diluted in fresh LB medium, containing 1 mg/mL mucin when required, and enumeration of CFU and PFU was performed. For the CFU enumeration, 1 mL aliquots of bacterial cultures were serially diluted in the saline buffer then platted on LB-agar plates (100 μL) and incubated overnight. For the PFU assessment, 1 mL aliquots were centrifuged 5 min at maximum speed (14000 rpm) for the collection of phage particles in the supernatant. Serial dilutions of these PFU were performed in SM buffer (100 mM NaCl, 10 mM MgSO4, 20 mM Tris-HCl, pH 7.5), then 10 μL of the pertinent dilutions were plated by the double-layer method (37). Two flasks per condition were considered as biological duplicates.

### Assessment of phage resistance

#### a. Spot test

The spot test assay was undertaken as described by Raya *et al*. (38). We used vB_KpnS-VAC35 WT, vB_KpnS-VAC35_ad15 and vB_KpnS-VAC35_ad15_m phages, that is prior to co-evolution, and adapted to K3574 during 15 days in the absence and presence of mucin, respectively.

#### b. Calculation of the frequency of phage-resistant mutants

The frequency of resistant mutants was calculated as previously described by Lopes *et al.* (39). Overnight cultures of the strains K3574 and K3325 at the different conditions evaluated were diluted 1:100 in LB and grown to an OD_600nm_ of 0.7. An aliquot of 1 mL of the culture containing 10^8^ CFU/mL was serially diluted, and the corresponding dilutions were mixed with 100 μL of vB_KpnS- VAC35 at 10^9^ PFU/mL, then plated by the double-layer method in TA medium. The plates were incubated at 37°C for 24h, then the colonies of resistant mutants were enumerated. The mutation rate was calculated by dividing the number of resistant bacteria (growing in the presence of the phage) by the total number of bacteria plated in conventional LB-agar (100 μL).

### Genomic DNA extraction and whole-genome sequencing

The DNeasy Blood & Tissue Kit (Qiagen®) was used for extracting the genomic DNA of bacterial cultures co-evolved 15 dpi with the vB_KpnS-VAC35 alone (K3574_ad15_P), in the presence of mucin (K3574_ad15_P+M) and exposed 15 days to mucin (K3574_ad15_M), following the manufacturer’s instructions. Samples were quantified with a Qubit 3.0 fluorometer using a Qubit dsDNA HS Assay Kit and with a Nanodrop spectrophotometer to evaluate the DNA purity. The MinION MK1C instrument with the Rapid Barcoding Kit (SQK-RBK004) were employed, following the manufacturer’s protocol and using a MinION flow cell v.9.4.1.

### Bioinformatic analysis

Basecalling was performed using GUPPY (Version 5.0.7 Super-accuracy model (SUP)) to generate fastQ sequencing reads from electrical data (the fast5 files generated by MinION). The reads were then further subsampled according to their barcodes and *de novo* assembled using Unicycler. Further analyses were performed after visualization of circular assembled genomes using Bandage. Draft assemblies were corrected by using iterative rounds of polishing with the Racon error correction software. Annotations were performed using Prokka (40), and insertions, deletions and other SNPs were called using the structural variant caller Snippy (v1.0.11). The presence and absence of intact genetic sequences were analyzed using Roary and Orthovenn2. OrthoVenn2 (http://www.bioinfogenome.net/OrthoVenn/) was used to compare the proteins of the four complete genomes using the files generated by Prokka analysis. Fasta files obtained after annotation were surveilled for indels using the blastn and blastp tools from the NCBI and compared to the reference genome (for K3574_WT, BioSample code SAMEA3649560 included in the European BioProject PRJEB10018). The workflow taken from Nanopore sequencing to the genomic analysis is summarized in Figure 5 a.

## Transparency declarations

The authors declare not to have conflict of interest

## References

1. Kamruzzaman M, Iredell JR. 2019. CRISPR-Cas System in Antibiotic Resistance Plasmids in *Klebsiella pneumoniae*. Front Microbiol 10:2934.

2. Mackow NA, Shen J, Adnan M, Khan AS, Fries BC, Diago-Navarro E. 2019. CRISPR-Cas influences the acquisition of antibiotic resistance in *Klebsiella pneumoniae*. PLoS One 14:e0225131.

3. Pacios O, Fernández-García L, Bleriot I, Blasco L, González-Bardanca M, López M, Fernández-Cuenca F, Oteo J, Pascual Á, Martínez-Martínez L, Domingo-Calap P, Bou G, Tomás M, (SEIMC) SGoMoAaRtAGobotSSoIDaCM. 2021. Enhanced Antibacterial Activity of Repurposed Mitomycin C and Imipenem in Combination with the Lytic Phage vB_KpnM-VAC13 against Clinical Isolates of *Klebsiella pneumoniae*. Antimicrob Agents Chemother 65:e0090021.

4. Majkowska-Skrobek G, Markwitz P, Sosnowska E, Lood C, Lavigne R, Drulis-Kawa Z. 2021. The evolutionary trade-offs in phage-resistant *Klebsiella pneumoniae* entail cross-phage sensitization and loss of multidrug resistance. Environ Microbiol.

5. Pacios O, Blasco L, Bleriot I, Fernandez-Garcia L, Gonzalez Bardanca M, Ambroa A, Lopez M, Bou G, Tomas M. 2020. Strategies to Combat Multidrug-Resistant and Persistent Infectious Diseases. Antibiotics (Basel) 9.

6. Uyttebroek S, Chen B, Onsea J, Ruythooren F, Debaveye Y, Devolder D, Spriet I, Depypere M, Wagemans J, Lavigne R, Pirnay JP, Merabishvili M, De Munter P, Peetermans WE, Dupont L, Van Gerven L, Metsemakers WJ. 2022. Safety and efficacy of phage therapy in difficult-to-treat infections: a systematic review. Lancet Infect Dis 22:e208–e220.

7. Denes T, den Bakker HC, Tokman JI, Guldimann C, Wiedmann M. 2015. Selection and Characterization of Phage-Resistant Mutant Strains of *Listeria monocytogenes* Reveal Host Genes Linked to Phage Adsorption. Appl Environ Microbiol 81:4295–305.

8. Lopatina A, Tal N, Sorek R. 2020. Abortive Infection: Bacterial Suicide as an Antiviral Immune Strategy. Annu Rev Virol 7:371–384.

9. Castillo JA, Secaira-Morocho H, Maldonado S, Sarmiento KN. 2020. Diversity and Evolutionary Dynamics of Antiphage Defense Systems in *Ralstonia solanacearum* Species Complex. Front Microbiol 11:961.

10. Ambroa A, Blasco L, Lopez M, Pacios O, Bleriot I, Fernandez-Garcia L, Gonzalez de Aledo M, Ortiz-Cartagena C, Millard A, Tomas M. 2021. Genomic Analysis of Molecular Bacterial Mechanisms of Resistance to Phage Infection. Front Microbiol 12:784949.

11. Schooley RT, Biswas B, Gill JJ, Hernandez-Morales A, Lancaster J, Lessor L, Barr JJ, Reed SL, Rohwer F, Benler S, Segall AM, Taplitz R, Smith DM, Kerr K, Kumaraswamy M, Nizet V, Lin L, McCauley MD, Strathdee SA, Benson CA, Pope RK, Leroux BM, Picel AC, Mateczun AJ, Cilwa KE, Regeimbal JM, Estrella LA, Wolfe DM, Henry MS, Quinones J, Salka S, Bishop-Lilly KA, Young R, Hamilton T. 2017. Development and Use of Personalized Bacteriophage-Based Therapeutic Cocktails To Treat a Patient with a Disseminated Resistant *Acinetobacter baumannii* Infection. Antimicrob Agents Chemother 61.

12. Law N, Logan C, Yung G, Furr CL, Lehman SM, Morales S, Rosas F, Gaidamaka A, Bilinsky I, Grint P, Schooley RT, Aslam S. 2019. Successful adjunctive use of bacteriophage therapy for treatment of multidrug-resistant *Pseudomonas aeruginosa* infection in a cystic fibrosis patient. Infection 47:665–668.

13. Dedrick RM, Guerrero-Bustamante CA, Garlena RA, Russell DA, Ford K, Harris K, Gilmour KC, Soothill J, Jacobs-Sera D, Schooley RT, Hatfull GF, Spencer H. 2019. Engineered bacteriophages for treatment of a patient with a disseminated drug- resistant *Mycobacterium abscessus*. Nat Med 25:730–733.

14. Riquelme SA, Ahn D, Prince A. 2018. *Pseudomonas aeruginosa* and *Klebsiella pneumoniae* Adaptation to Innate Immune Clearance Mechanisms in the Lung. J Innate Immun 10:442–454.

15. Pletzer D, Mansour SC, Wuerth K, Rahanjam N, Hancock RE. 2017. New Mouse Model for Chronic Infections by Gram-Negative Bacteria Enabling the Study of Anti-Infective Efficacy and Host-Microbe Interactions. mBio 8.

16. Leão RS, Pereira RH, Folescu TW, Albano RM, Santos EA, Junior LG, Marques EA. 2011. KPC-2 carbapenemase-producing *Klebsiella pneumoniae* isolates from patients with Cystic Fibrosis. J Cyst Fibros 10:140–2.

17. Delfino E, Giacobbe DR, Del Bono V, Coppo E, Marchese A, Manno G, Morelli P, Minicucci L, Viscoli C. 2015. First report of chronic pulmonary infection by KPC-3- producing and colistin-resistant *Klebsiella pneumoniae* sequence type 258 (ST258) in an adult patient with cystic fibrosis. J Clin Microbiol 53:1442–4.

18. Moulton-Brown CE, Friman VP. 2018. Rapid evolution of generalized resistance mechanisms can constrain the efficacy of phage-antibiotic treatments. Evol Appl 11:1630–1641.

19. Hansson GC. 2019. Mucus and mucins in diseases of the intestinal and respiratory tracts. J Intern Med 285:479–490.

20. Paone P, Cani PD. 2020. Mucus barrier, mucins and gut microbiota: the expected slimy partners? Gut 69:2232–2243.

21. Carroll-Portillo A, Lin HC. 2021. Exploring Mucin as Adjunct to Phage Therapy. Microorganisms 9.

22. Jass JR, Walsh MD. 2001. Altered mucin expression in the gastrointestinal tract: a review. J Cell Mol Med 5:327–51.

23. de Freitas Almeida GM, Hoikkala V, Ravantti J, Rantanen N, Sundberg LR. 2022. Mucin induces CRISPR-Cas defense in an opportunistic pathogen. Nat Commun 13:3653.

24. Green SI, Gu Liu C, Yu X, Gibson S, Salmen W, Rajan A, Carter HE, Clark JR, Song X, Ramig RF, Trautner BW, Kaplan HB, Maresso AW. 2021. Targeting of Mammalian Glycans Enhances Phage Predation in the Gastrointestinal Tract. mBio 12.

25. Oechslin F. 2018. Resistance Development to Bacteriophages Occurring during Bacteriophage Therapy. Viruses 10.

26. Rendueles O, de Sousa JAM, Rocha EPC. 2023. Competition between lysogenic and sensitive bacteria is determined by the fitness costs of the different emerging phage- resistance strategies. Elife 12.

27. Barr JJ, Auro R, Furlan M, Whiteson KL, Erb ML, Pogliano J, Stotland A, Wolkowicz R, Cutting AS, Doran KS, Salamon P, Youle M, Rohwer F. 2013. Bacteriophage adhering to mucus provide a non-host-derived immunity. Proc Natl Acad Sci U S A 110:10771–6.

28. Almeida GMF, Laanto E, Ashrafi R, Sundberg LR. 2019. Bacteriophage Adherence to Mucus Mediates Preventive Protection against Pathogenic Bacteria. mBio 10.

29. Wright RCT, Friman VP, Smith MCM, Brockhurst MA. 2019. Resistance Evolution against Phage Combinations Depends on the Timing and Order of Exposure. mBio 10.

30. Ohshima Y, Schumacher-Perdreau F, Peters G, Pulverer G. 1988. The role of capsule as a barrier to bacteriophage adsorption in an encapsulated *Staphylococcus simulans* strain. Med Microbiol Immunol 177:229–33.

31. Hao G, Shu R, Ding L, Chen X, Miao Y, Wu J, Zhou H, Wang H. 2021. Bacteriophage SRD2021 Recognizing Capsular Polysaccharide Shows Therapeutic Potential in Serotype K47 *Klebsiella pneumoniae* Infections. Antibiotics (Basel) 10.

32. Blasco L, López-Hernández I, Rodríguez-Fernández M, Pérez-Florido J, Casimiro- Soriguer CS, Djebara S, Merabishvili M, Pirnay J-P, Rodríguez-Baño J, Tomás M, López Cortés LE. 2023. Case report: Analysis of phage therapy failure in a patient with a *Pseudomonas aeruginosa* prosthetic vascular graft infection. Frontiers in Medicine 10.

33. Liu M, Hernandez-Morales A, Clark J, Le T, Biswas B, Bishop-Lilly KA, Henry M, Quinones J, Voegtly LJ, Cer RZ, Hamilton T, Schooley RT, Salka S, Young R, Gill JJ. 2022. Comparative genomics of Acinetobacter baumannii and therapeutic bacteriophages from a patient undergoing phage therapy. Nat Commun 13:3776.

34. Deutscher J, Francke C, Postma PW. 2006. How phosphotransferase system-related protein phosphorylation regulates carbohydrate metabolism in bacteria. Microbiol Mol Biol Rev 70:939–1031.

35. Bergstrom K, Shan X, Casero D, Batushansky A, Lagishetty V, Jacobs JP, Hoover C, Kondo Y, Shao B, Gao L, Zandberg W, Noyovitz B, McDaniel JM, Gibson DL, Pakpour S, Kazemian N, McGee S, Houchen CW, Rao CV, Griffin TM, Sonnenburg JL, McEver RP, Braun J, Xia L. 2020. Proximal colon-derived O-glycosylated mucus encapsulates and modulates the microbiota. Science 370:467–472.

36. Mangalea MR, Duerkop BA. 2020. Fitness Trade-Offs Resulting from Bacteriophage Resistance Potentiate Synergistic Antibacterial Strategies. Infect Immun 88.

37. Abedon ST, Yin J. 2009. Bacteriophage plaques: theory and analysis. Methods Mol Biol 501:161–74.

38. Raya RR, H’bert EM. 2009. Isolation of Phage via Induction of Lysogens. Methods Mol Biol 501:23–32.

39. Lopes A, Pereira C, Almeida A. 2018. Sequential Combined Effect of Phages and Antibiotics on the Inactivation of *Escherichia coli*. Microorganisms 6.

40. Seemann T. 2014. Prokka: rapid prokaryotic genome annotation. Bioinformatics 30:2068–9.

